# B Cell Receptor Signaling Drives APOBEC3 Expression Via Direct Enhancer Regulation in Chronic Lymphocytic Leukemia B Cells

**DOI:** 10.1101/2021.07.27.454050

**Authors:** Zhiquan Wang, Huihuang Yan, Justin C. Boysen, Charla R. Secreto, Jian Zhong, Jiaqi Zhou, Haiyun Gan, Chuanhe Yu, Esteban Braggio, Susan L. Slager, Sameer A. Parikh, Neil E. Kay

## Abstract

Constitutively activated B cell receptor (BCR) signaling is a primary biological feature of chronic lymphocytic leukemia (CLL). The biological events controlled by BCR signaling in CLL are not fully understood and need investigation. To make inroads we obtained blood samples from CLL patients before and after Bruton’s tyrosine kinase inhibitors (BTKi) treatment and used them to study BCR signaling regulated genes. Here, by analysis of the chromatin states and gene expression profiles of CLL B cells from patients before and after BTKi ibrutinib treatment, we show that BTKi treatment leads to a decreased expression of APOBEC3 family genes in an enhancer regulation dependent manner. BTKi treatment reduces enrichment of enhancer markers (H3K4me1, H3K27ac) and chromatin accessibility at putative APOBEC3 enhancers. CRISPR-Cas9 directed deletion or inhibition of the putative APOBEC3 enhancers leads to reduced APOBEC3 expression. We further find that transcription factor NFATc1 couples BCR signaling with the APOBEC3 enhancer activity to control APOBEC3 expression. Importantly, enhancer regulated APOBEC3 expression contributes to replication stress in malignant B cells. We also demonstrate a novel mechanism for BTKi suppression of APOBEC3 expression via direct enhancer regulation in a NFATc1 dependent manner, implicating BCR signaling as a potential regulator of leukemic genomic instability.

**Key points:**

- BCR signaling pathway regulates APOBEC3 expression via direct enhancer regulation.
- AOPEBC3 enhancers are involved in the process of DNA replication stress, implicating a potential role in B cell genomic instability and CLL evolution

## Introduction

CLL is the most common leukemia in the U.S. with ~21,000 new cases diagnosed each year and characterized by constitutively activated BCR signaling pathway^1^. BCR signaling has a crucial role both in normal B cell development and B cell malignancies. During normal development, B cells are derived from the bone marrow hematopoietic stem cells and maturate through the expression of a functional BCR. Experienced B cells can specifically recognize foreign antigens via their unique BCRs, which in turn triggers antigen specific antibody responses and promotes B cell differentiation into plasma cells and memory B cells^2^. In CLL, the BCR signaling pathway may be activated by antigens in the tissue microenvironment or occur because of mutations in the BCR signaling genes to promote leukemic cell maintenance and expansion^1,3^. Active BCR signaling is mediated through activation of downstream kinases, such as spleen tyrosine kinase (SYK), Bruton tyrosine kinase (BTK) and Phosphoinositide 3-kinases (PI3K)^2^. These kinases have become key therapeutic targets to inhibit the BCR signaling for the treatment of B cell malignancies. Indeed, BTKi inhibitors (BTKis) such as ibrutinib, acalabrutinib, zanabrutinib have remarkable efficacy in the treatment of patients with CLL^4–6^. BTKi treatment can suppress expression of genes related to pathways including BCR signaling, NF-κB signaling pathways, inflammation, cytokine signaling, cell adhesion, p53 response, MAPK signaling, cell cycle, focal adhesion, calcium signaling, and Wnt signaling ^7,8,9^. However, there is still a lack of understanding how BCR signaling regulates downstream gene expression in CLL. Therefore, exploring the mechanism whereby BTKi regulates downstream gene expression would provide insights into the understanding of BCR signaling pathway as well as new directions for CLL treatment.

Cell signaling pathways can regulate gene expression through modification of the epigenetic states of cells^10–12^. Recently, epigenomic surveys in CLL detected alterations of epigenetic landscapes as well as mutations of genes encoding key chromatin machineries^13–15^. BTKi treatment can lead to epigenetic reprogramming of patients CLL B cells^9^. These findings suggest that epigenetic events are involved in the BCR signaling pathway. However, the mechanisms and functional importance of these epigenetic programs in CLL and BCR signaling are largely unknown.

One major mechanism that epigenetic programs utilize to control gene expression and cell states is to regulate the activity of enhancers, a class of regulatory DNA elements capable of stimulating transcription over long genomic distances^16^. At enhancers, transcription factors (TFs) trigger the recruitment of chromatin-modifying enzymes to establish active histone modifications on adjacent nucleosomes, such as histone H3 lysine 27 acetylation (H3K27ac) and histone H3 lysine 4 mono-methylation (H3K4me1)^17^. As a result, the active enhancers can promote their target genes expression and related cellular functions. Therefore, genome-wide measurements of H3K27ac and H3K4me1 enrichment (e.g., using chromatin immunoprecipitation [ChIP-seq] or CUT&Tag) and chromatin accessibility (e.g., using the assay for transposase-accessible chromatin [ATAC-seq]) allow for the annotation of active enhancer landscapes in different cellular states^18,19^.

Here, by characterization of the gene expression and enhancer profiles of CLL B cells from patients before and while on ibrutinib treatment, we found that BCR signaling pathway drives Apolipoprotein B mRNA-editing enzyme, catalytic polypeptide-like type 3 (APOBEC3) expression via NFATc1-dependent enhancer regulation. We also showed that AOPEBC3 enhancer is involved in the process of DNA replication stress, implicating a potential role in B cell genomic instability and CLL evolution.

## Results

### BCR signaling regulates APOBEC3 expression in CLL

To explore BCR signaling regulated genes in CLL, we analyzed the gene expression profile of CLL B cells from patients before and after ibrutinib treatment by mRNA-seq (n = 8 patients, at around one-year of continuous ibrutinib treatment, **Supplementary table 1**). BTKi treatment induced dramatic gene expression changes (total changed genes = 3334, up =1964, down=1370. p < 0.05, fold change > 1.5) (**Supplementary Fig 1A, Supplementary table 2**). As reported before, BTKi suppresses the gene expression involved in mitochondrial function^20^ and BCR signaling pathway (**Fig 1A, Supplementary table 3, Supplementary Fig 1 B**). We also found other known BTKi regulated pathways through gene ontology pathway analysis (**Supplementary table 4**). In addition to these pathways involved in B cell malignancies and CLL survival, we found that ibrutinib treatment led to the reduction of expression in genes associated with single strand DNA deamination (**Fig 1A, B, C, Supplementary Fig. 1C**). The BTKi regulated genes involved in this process mainly contains the APOBEC3 family genes (*APOBEC3C, APOBEC3D APOBEC3F, APOBEC3G, APOBEC3H*)^21^, and their expression levels showed a consistent reduction in CLL B cells from ibrutinib treated patients (**Fig. 1D**). Further analysis of published single cell RNA-seq data^9^ revealed that BTKi treatment downregulated the expression of APOBEC3C in CLL B cells from one patient and APOBEC3G in CLL B cells of all three patients (**Fig. 1E, Supplementary Fig. 1D, E**). We then confirmed the reduction of APOBEC3 levels by RT-qPCR and western blot in CLL B cells from four patients before and after one-year of continuous ibrutinib treatment (**Supplementary Fig. 1F and Fig. 1F, G**). Treatment of purified primary CLL B cells isolated from ibrutinib untreated patients and cultured with ibrutinib at a sublethal level could also reduce the APOBEC3 expression, indicating the reduced expression of these genes in patients is not due to the elimination of APOBEC3 high expressed cells (**Supplementary Fig. 1G**). We also compared the basal expression of APOBEC3 between CLL B cell and normal B cells using data from public datasets ^22,23^ and found an increased expression of APOBEC3 in CLL B cells versus the normal B cells (**Supplementary Fig. 2A, B**). We then confirmed the increased expression of APOBEC3C and APOBEC3G by western blot in CLL B cells compared to normal B cells (**Supplementary Fig. 2C, D**). Together, our results indicated that active BCR signaling may promote APOBEC3 expression in leukemic cells of CLL patients.

**Figure 1.**
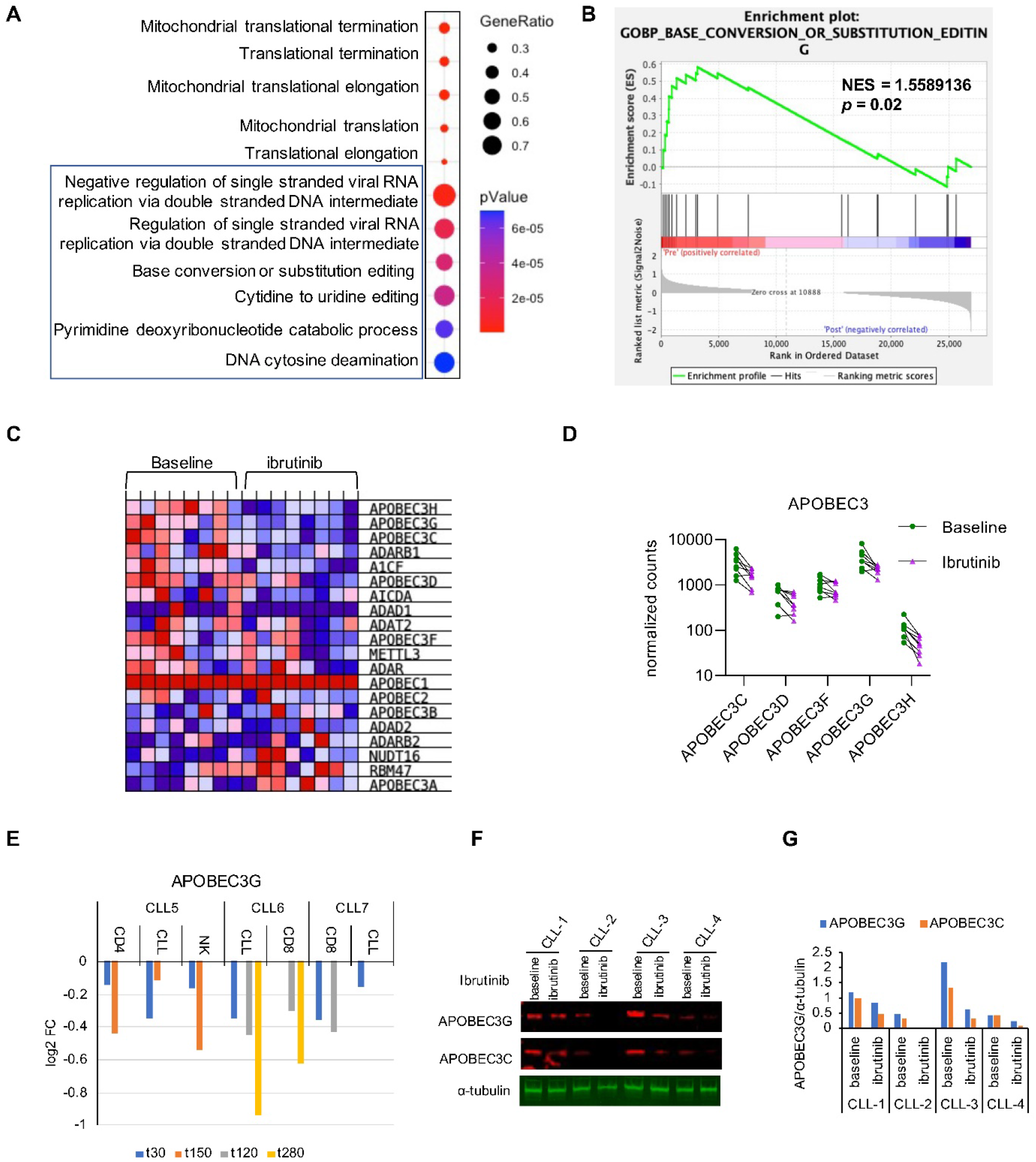
BCR signaling dependent APOBEC3 expression in CLL B cells. A. Gene Ontology (GO) enrichment analysis for the ibrutinib suppressed genes. CLL B cells were harvested from the same CLL patients before and with one-year continuous ibrutinib treatment and the gene expression was determined by RNA-seq, n=8 patients.
B. Gene set enrichment analysis showing enrichment of base conversion or substitution editing linked genes in ibrutinib pretreated CLL B cells compared to CLL B cells from patients with continuous one-year ibrutinib treatment.
C. Blue-Pink O’Gram in the Space of the Analyzed GeneSet in panel B). Red and blue color indicated higher and lower expression respectively.
D. APOBEC3 genes expression change in CLL B cells from ibrutinib treated patients compared to that of pretreated patients. baseline = pre ibrutinib treatment; ibrutinib = one-year of continuous ibrutinib treatment.
E. APOBEC3G expression in indicated cells from ibrutinib treated patients. t30 means 30 days ibrutinib treatment.
F. Immunoblot analysis of APOBEC3C and APOBEC3G in CLL B cells from patients before and after one-year ibrutinib treatment. baseline = pre ibrutinib treatment; ibrutinib = one-year ibrutinib treatment.
G. Quantification of the western blot intensity in panel F.

### BTKi treatment suppresses the activity of putative enhancers of APOBEC3

Signaling pathways can regulate chromatin modifying enzymes, histone modifications, and nucleosome occupancy to affect both epigenetic and transcriptional state of cells^10,11^. APOBEC3 are genes clustered in tandem on chromosome 22^24^. We hypothesized that BCR signaling regulates APOBEC3 expression by modifying the local chromatins around the APOBEC3 gene cluster. To check the local chromatin states of APOBEC3, we performed CUT&Tag^25^ to map the histone marks including H3K4me1, H3K4me3, H3K27ac and ATAC-seq to examine the chromatin accessibility of the leukemic cells from CLL patients before and with one-year of continuous ibrutinib treatment (**Fig. 2A, Supplementary table S1**). We found the enrichment of enhancer marker H3K4me1 proximal to the APOBEC3 gene clusters (chr22:39,060,747-39,101,866), and these regions were also enriched with active enhancer marker H3K27ac and had an open chromatin state (Fig 2B, Supplementary Fig. 3). This suggests that this region is a putative enhancer(s) that controls APOBEC3 expression. Hereafter we refer this region as APOBEC3 enhancers (AEs). We further found that BTKi treatment caused reductions of H3K4me1, H3K27ac, and chromatin accessibility at these regions in most ibrutinib treated patients, however, there was no change of the promoter marker H3K4me3 in these genes (**Fig 2B, Supplementary Fig 3**), which indicated that BTKi treatment leads to APOBEC3 genes expression change via the regulation of their enhancer regulation. Next, we analyzed the published ATAC-seq data of CLL B cells from patients who are on ibrutinib treatment^9^. While the ibrutinib treatment in this study was shorter than ours (120 days after treatment), it did lead to modest reductions of chromatin accessibility of AEs in most patients (**Supplementary Fig. 4**). Thus, these results indicate that BTKi may induce decreased expression of APOBEC3 genes through regulation of the activity of their enhancers.

**Figure 2.**
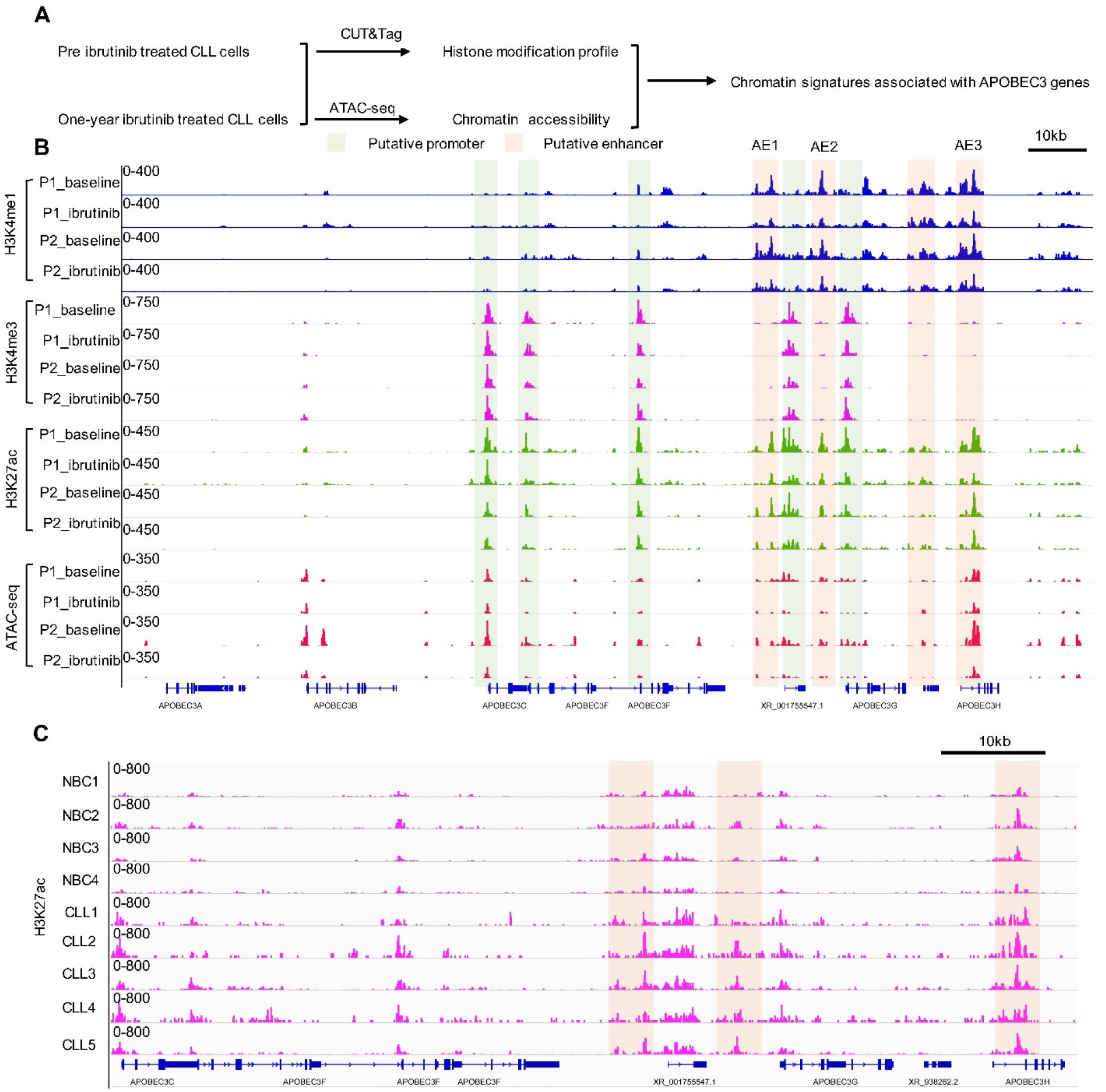
Ibrutinib treatment suppresses the activity of putative enhancers of APOBEC3. A. Schematic view of the analysis of epigenetic signatures of CLL B cells with one-year of continuous ibrutinib treatment.
B. Genome tracks showing CUT&Tag of H3K4me1, H3K4me3, H3K27ac, and ATAC-seq profiles of putative APOBEC3 enhancers. Light green and brown shadows show promoters and enhancers respectively. P, patient; baseline = pre ibrutinib treatment; ibrutinib = one-year of continuous ibrutinib treatment.
C. Genome tracks showing H3K27ac profiles of APOBEC3 enhancers from CUT&Tag on normal and CLL B cells.

Given this we further analyzed the published datasets^23^ to compare the enhancer signatures between CLL B cells and normal B cells. CLL B cells showed higher enrichment of H3K27ac, and increased chromatin accessibility at AEs (**Supplementary Fig. 5A**). Then we performed the H3K27ac CUT&Tag in normal B cells and CLL B cells and our results also showed that CLL B cells had higher H3K27ac at the AE regions (**Fig. 2C**). Analyzing the Encode H3K27ac ChIP-seq data found that this enhancer character is limited to the B cell lineages in hematopoietic cell populations (**Supplementary Fig. 5B**). Importantly, the Hi-C^26^ data generated from B-lymphocyte cell line GM12878 cells showed that the AOPBEC3 genes and enhancers had high level of genomic interactions and located in the same topologically associating domain (TAD) (**Supplementary Fig 5C**). Together, our results indicated that the expression of APOBEC3 may be controlled by BCR signaling through enhancer regulation and modifications.

### APOBEC3 expression is controlled by their enhancer activity

Based on the enrichment of H3K4me1, H3K27ac and chromatin accessibility, AE regions contains three active enhancer modules, we designated these modules as AE1, AE2, and AE3(**Fig. 2B, Fig. 3A**). To assess the functional activity of these enhancers on the expression of APOBEC3 genes, we investigated the consequence of deletion of each one of these AEs in MEC1 cell line. MEC1 was used based on fact that MEC1 has been utilized in previous CLL epigenetic study^27^ and we also found that it has similar chromatin states at the AEs, though the relative enrichment of enhancer marks (H3K4me1, H3K27ac and ATAC-seq intensity) at AE1 and AE2 is lower compared to that of the primary CLL B cells (**Fig. 4F and Supplementary Fig. 6G**). Initially we tried to generate AE knockout clones by CRISPR mediated knockout, however, we failed to grow cells from single clones with sgRNA transfected MEC1 cells (data not shown). Next, we infected the Cas9 expressing MEC1 cells with sgRNAs to generate pool populations of cells with AEs deletion (**Fig 3A**). PCR analysis showed very robust deletion of AE1, AE2, and AE3 (**Fig 3B**). Both deletion of AE1 or AE2 reduced the expression of APOBEC3 genes (**Fig. 3C, D**), while AE3 deletion suppressed the expression of APBEC3C, APOBEC3D, APOBEC3F and APOBEC3G, but not APOBEC3H, which is closest to AE3 (**Fig. 3C, D**). Overall, AE2 deletion led to the strongest decrease of all the APOBEC3 genes.

**Figure 3.**
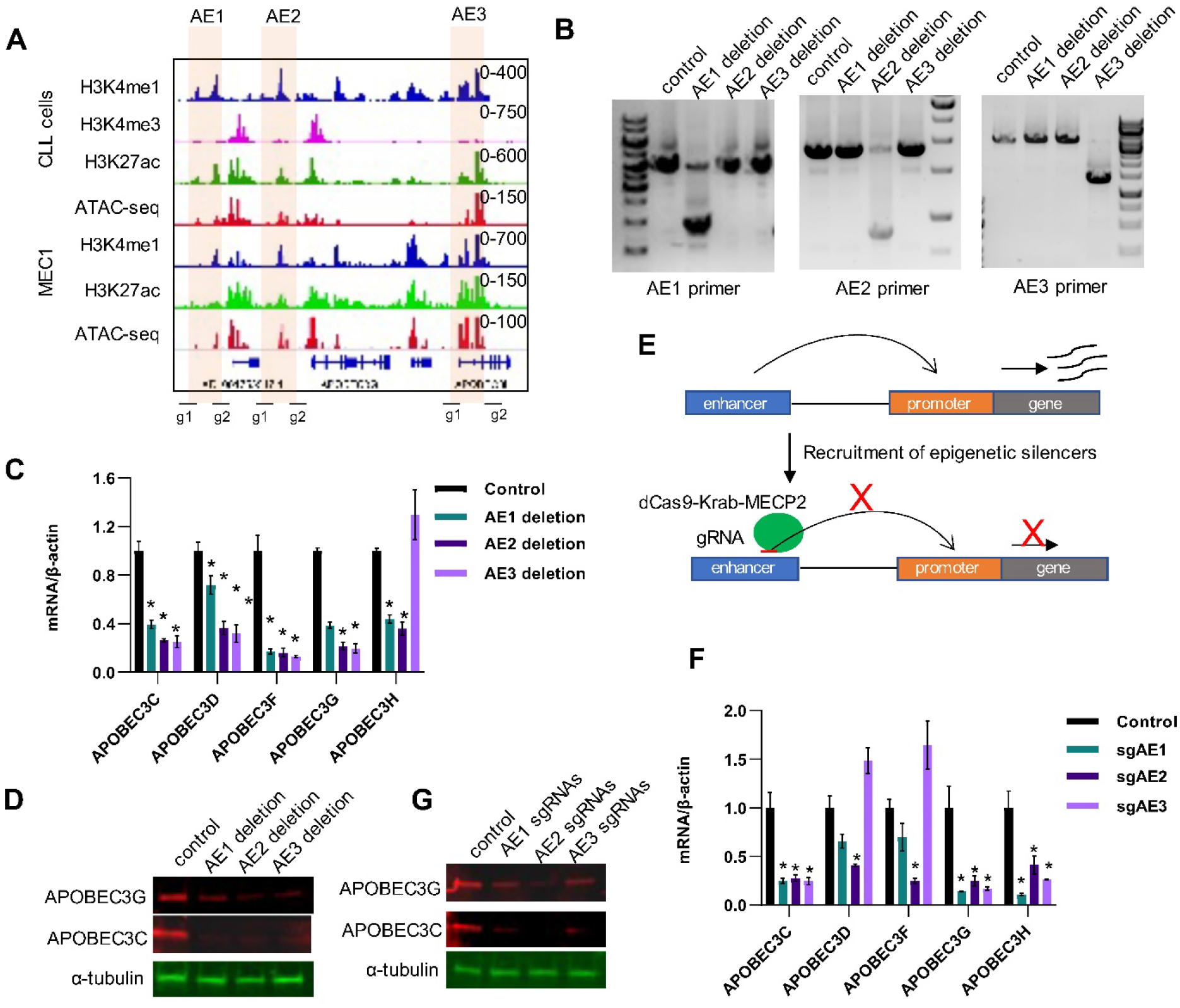
Deletion or inhibition of APOBEC3 enhancers suppresses APOBEC3 expression. A. An outline of the genomic locus targeted by the gRNAs to the AE enhancers.
B. Gel imaging showing successful deletion of AE enhancers. The AE enhancer regions were amplified by PCR primers spanning the indicated regions.
C. RT-qPCR analysis of APOBEC3 expression in the AE deleted MEC1 cells. n = 3 independent experiments.
D. Western blot analysis of APOBEC3C and APOBEC3G expression in the AEs deleted MEC1 cells.
E. Schematic view of the strategy of CRISPRi.
F. RT-qPCR analysis of APOBEC3 expression after inhibition of individual AEs by CRISPRi. n = 3 independent experiments.
G. Western blot analysis of APOBEC3C and APOBEC3G expression after inhibition of individual AEs by CRISPRi in MEC1 cells.

**Figure 4.**
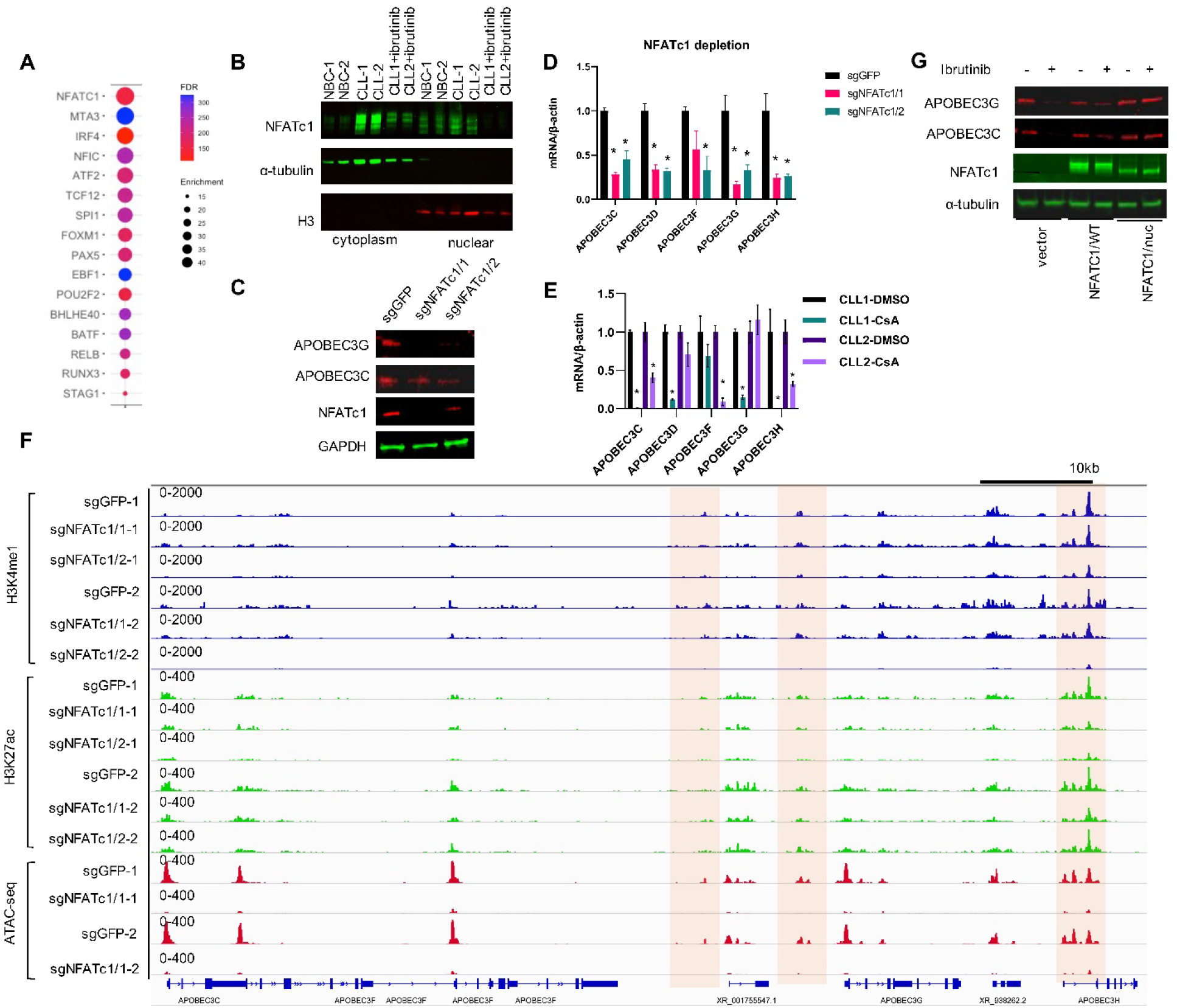
NFATc1 controls the APOBEC3 expression through enhancer regulation. A. Top TF motifs enriched in the regions with decreased chromatin accessibility in CLL B cells after one-year of continuous ibrutinib treatment are shown.
B. Western blot showing protein levels of NFATc1 in nuclear and cytoplasm fraction of CLL B cells from patients treated with or without ibrutinib.
C. Western blot showing protein levels as indicated in NFATc1 depleted and control MEC1 cells.
D. RT-qPCR analysis of APOBEC3 expression in NFATc1 depleted MEC1 cells. n = 3 independent experiments.
E. RT-qPCR analysis of APOBEC3 expression in CLL B cells treated with 2.5 μM Cyclosporin A for 24 hours. n = 3 independent experiments for each CLL sample.
F. Genome tracks showing CUT&Tag of H3K4me1, H3K4me3, H3K27ac, and ATAC-seq profiles of APOBEC3 genes in NFATc1 depleted and control MEC1 cells.
G. Western blot showing protein levels as indicated in wildtype (NFATc1/wt) or nuclear stable form (NFATc1/nuc) NFATc1 expressing MEC1 cells treated with or without ibrutinib. NFAT1c1 was detected by Flag antibody. Cells were treated with ibrutinib at 2.5μM for 3 days.

Cas9 introduces DNA breaks at a specified site to inactivate gene function but the reliance on endogenous DNA repair machinery can make it difficult to limit the outcome to a single desired change^28^. A recent study has shown that CRIPSR Cas9 can even induce chromothripsis as an on-target consequence^29^. To exclude that the observed results with AE deletions above were outcomes of undesired changes induced by Cas9, we used CRISPR interference (CRISPRi) to modulate enhancer activity by rewriting the epigenetic states without changing the underlying DNA sequence.^30,31^. CRISPRi induces a suppressive chromatin state at targeted loci through recruitment of a KRAB effector domain and a MECP2 fused to a catalytically dead Cas9 (dCas9-Krab-MECP2) (**Fig. 3E**). RT-PCR and western blot analysis showed a significant reduction of most of the APOBEC3 genes expression by gRNAs targeting all three AEs (**Fig 3F, G**), though there were some inconsistencies to the AE KO assays regarding the effect of targeting the AE3 on APOBEC3H expression (**Fig. 3C, F**). These inconsistencies might result from the complicated local chromatin structure that affect the process of Cas9-KRAB-MECP2 mediated epigenetic silencing^32^. Together, we identified the BCR signaling dependent enhancers that regulate APOBEC3 expression. All three enhancer modules (AE1, AE2, AE3) are able to regulate most of the APOBEC3 gene expression, these results are consistent with the recent report that individual elements of a super-enhancer could contribute to their target gene expression^33^.

### NFATc1 controls APOBEC3 enhancer activity

We reasoned that identifying transcription factors (TF) enriched at regions with altered chromatin states would allow us to determine the mechanism that links BCR signaling to epigenetic regulation of AE activity and APOBEC3 expression. To that end, we evaluated whether specific TF motifs were enriched within regions with chromatin accessibility reduction after ibrutinib treatment. The top enriched motifs included binding sites for the NFATc1, ATF2, and IRF4 (**Fig. 4A**). We focused on NFATc1, a transcription factor that can promote enhancer reprogramming^34^ and importantly it is known that, NFATc1 is a putative downstream factor of BCR in CLL^35,36^ and has been shown to regulate APOBEC3G expression^37,38^. Moreover, NFATc1 is also enriched at the APOBEC3 enhancer regions in the GM12878 cells (**Supplementary Fig. 6A**). Western blots showed a reduction of nuclear NFATc1 level in CLL B cells of ibrutinib treated patients (**Fig. 4B, Supplementary Fig. 6B**), suggesting that BTKi treatment depletes the nuclear fraction of NFATc1, which may in turn abolish or reduce its function. We found that, BTKi treatment led to a reduction of total NFATc1 both at a protein (**Fig. 4B**) and RNA level (**Supplementary Fig. 6C**) and an even more robust reduction of the nuclear fraction. We also observed decreased chromatin accessibility and H3K4me1, H3K27ac around the gene locus of NFATc1 in CLL B cells from BTKi treated patients (**Supplementary Fig. 6D**). Although NFATc2 binds to similar DNA motifs as NFATc1, we did not see changes of NFATc2 with ibrutinib treatment (**Supplementary Fig. 6C**). These results suggested the role of NFATc1 in the regulation of AE activity and APOBEC3 genes expression.

We next explored the function of NFATc1 in the regulation of APOBEC3 expression. Depletion of NFATc1 led to a reduced expression of APOBEC3 in MEC1 cells, whereas depletion of another top hit TF IRF4 had no effect on APOBEC3 expression (**Fig. 4C, D, Supplementary Fig. 6E, F**). Importantly, treating the primary CLL B cells with a Calcineurin inhibitor Cyclosporin A (CsA), which can suppress the NFATc1 nuclear localization^39^, suppressed the APOBEC3 expression (**Fig. 4E**). We next performed CUT&Tag and ATAC-seq in NFATc1 depleted and control MEC1 cells to test if NFATc1 is required for the active APOBEC3 enhancer to promote APBEC3 expression. NFATc1 depletion resulted in decreased chromatin accessibility and H3K27ac enrichment at AEs (**Fig. 4F, Supplementary Fig. 6G**), which demonstrated that NFATc1 is the key factor that maintains the active chromatin state of APOBEC3 enhancers.

To test our hypothesis that BTKi treatment depletes the nuclear NFATc1, which in turn abolishes the AEs activity and APOBEC3 expression, we overexpressed a nuclear stable form of NFATc1 (NFATc1^nuc^, this form of NFATc1 is constitutively located in the nucleus)^40^ in MEC1 cells and found that it could block the BTKi induced APOBEC3 reduction (**Fig 4G**). This result supports that nuclear depletion of NFATc1 is required for the BTKi induced APOBEC3 expression reduction. Here our results suggest that NFATc1 couples the epigenetic program and BCR signaling to control APOBEC3 expression.

### APOBEC3 enhancers contribute to DNA replication stress in CLL B cells

Next, we sought to determine the role of APOBEC3 enhancers in CLL B cells in relation to DNA stress as APOBEC3 enzymes have been implicated in somatic mutagenesis and cancer evolution^21^. Recently, APOBEC3 induction has been shown to increase DNA replication stress and chromosome instability in early breast and lung cancer evolution^41^. There are also reports showing that APOBEC family mutational signatures are associated with development and poor prognosis of multiple myeloma^42,43^. However, there is little understanding of the function of APOBEC3 in CLL B cells. While APOBEC3A and APOBEC3B are thought to be the main family members involved in APOBEC3 mediated cancer mutagenesis, we found APOBEC3C-H are the major family members upregulated by BCR signaling in CLL. Indeed, APOBEC3C, APOBEC3G, and APOBEC3H have been reported to potentially target host genomes^44,45^. Since APOBEC3 deaminates ssDNA, we reasoned that APOBEC3 in CLL B cells may also contribute to replication stress and DNA instability. We evaluated the function of APOBEC3 in MEC1 cells by deletion of the APOBEC3 enhancers or only AE2, which can downregulate the expression of most APOBEC3 genes (**Fig3**). We found that MEC1 cells have a high level of spontaneous DNA damage, illustrated by phosphorylated pChk1 (S345)^46^, 53BP1 nuclear body (a marker of DNA replication stress^47^), accumulation of RPA2 positive cells^48^ and DNA damage marker gamma H2Ax (γH2Ax) in the S phase cells^49^ (**Fig. 5A-D**). However, AE-deleted MEC1 cells showed a reduction of pChk1 (S345) compared to MEC1 control cells (**Fig. 5A**). In addition, compared to the control cells, AE2 deleted cells had fewer 53BP1 nuclear bodies (**Fig. 5B**) and RPA2 positive cells (**Fig. 5C**), both of which are associated with replication stress induced DNA damage response. Importantly, AE2 deleted cells also showed decreased γH2Ax in the S phase cells (**Fig. 5D**). We next performed Edu/PI assay to check the role of APOBEC3 in DNA replication. Consistently, we noticed that MEC1 cells had a fraction of S phase cells with low Edu incorporation during S phase, indicating DNA replication stress (**Fig. 5E**); however, AE2 deletion greatly increased Edu incorporation (**Fig 5E**). Taken together, these data suggest that increased expression of APOBEC3 may be involved in DNA replication stress and drives genomic instability in malignant B cells.

**Fig. 5.**
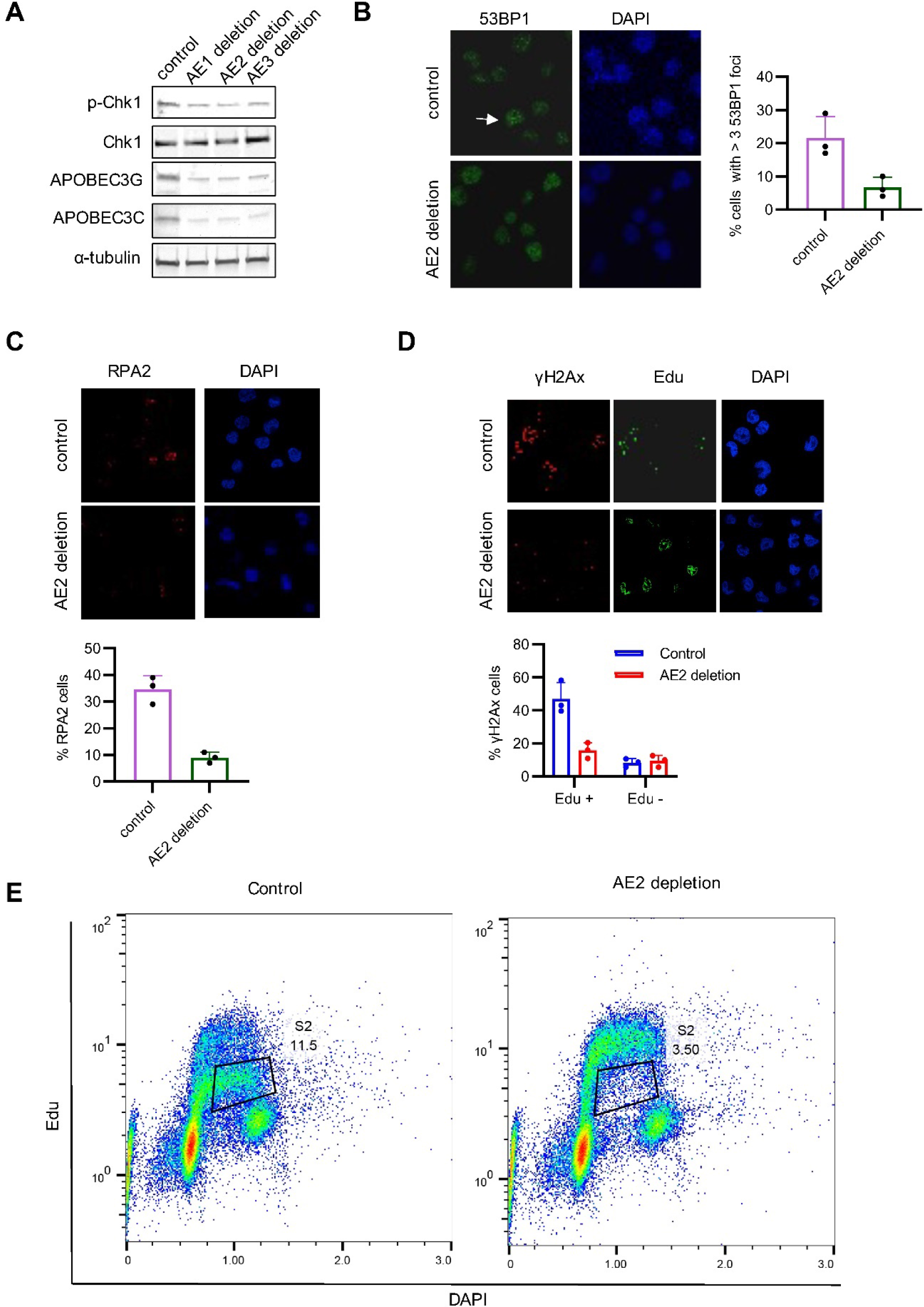
APOBEC3 enhancers contribute to replication stress induced DNA damage in MEC1 cells. A. Western blot analysis of pChk1 level in the indicated AE deleted MEC1 cells.
B. Representative images of 53BP1 staining in AE2 depleted and control MEC1 cells. Right panel shows the quantification of cells with more than 3 53BP1 foci (100 cells were counted per experiment, n = 3 independent experiments).
C. Representative images of RPA staining in AE2 depleted and control MEC1 cells. Right panel shows the quantification of cells with RPA2 (100 cells were counted per experiment, n = 3 independent experiments).
D. Representative images of γH2AX and Edu staining of AE2 depleted and control MEC1 cells treated. Cells were incubated with 10 μM Edu for 30 mins before harvest. (100 cells were counted per experiment, n = 3 independent experiments).
E. AE2 deleted and control MEC1 cells were incubated with Edu for 30min before the cells were stained with Click-it Alexa 488 azide and DAPI.

## Discussion

In recent years, the implement of highly specific, targeted novel agents for human malignancies has greatly improved the outcome of certain diseases. We have reasoned that patient derived tissues while on or even after novel agent targeted therapy can provide powerful tools to study the gene expression regulatory networks related to the targeted protein. Supporting this concept, we have explored the epigenome and transcriptome of CLL B cells from patients before and after the novel agent ibrutinib treatment and demonstrated that the BCR signaling pathway in leukemic B cells regulates APOBEC3 expression via direct regulation of the enhancer activity of the APOBEC3 gene cluster. In addition, this BCR mediated APOBEC3 gene expression is suppressed by a BTK inhibitor, ibrutinib. Overall, using ibrutinib treated CLL patients as a model, we demonstrate that this novel BTKi targeted agent can be used to gain important insights and provides a valuable resource in the study of basic biologic questions of B cell biology.

Signaling pathways can regulate gene expression through modification of the epigenetic states of the cells^10,11^, and most knowledge is gained from stem cells studies ^50,51^. This mechanism is also utilized by oncogenic signaling pathways to activate transcription of genes that promote malignant cell survival and growth^52,53^. Here our results provide evidence that a signaling pathway which regulates gene expression through epigenetic modification, exists in leukemic CLL B cells. We showed that NFATc1 is activated by the BCR signaling pathway, which in turn activates the enhancer activity of the APOBEC3 genes to promote their expression. Although NFATc1 has been shown to be involved in the regulation of Epstein Barr virus (EBV) associated super-enhancers, the exact mechanism by which NFATc1 controls the activity of enhancers in CLL unknown. Previous studies have shown that NFATc1 binds to several chromatin regulators (e.g., p300)^54^, so it is possible that NFATc1 works to recruit the chromatin modification enzymes and remodelers to these regions to generate an open chromatin structure for APOBEC3 expression. Indeed, our data shows that NFATc1 depletion leads to decreased chromatin accessibility and active histone modifications at the AEs. Future studies to explore the chromatin regulators that work with NFATc1 to regulate the chromatin states could provide more information about this regulatory aspect in the mechanism of APOBEC3 expression. Despite our findings that BTKi treatment suppresses APOBEC3 expression in an enhancer dependent manner, it remains unclear what is the biological function of this process. APOBEC3 family members play important roles in intrinsic responses to infection by retroviruses and have been implicated in the control of other viruses, such as parvoviruses, herpesviruses, papillomaviruses, hepatitis B virus, and retrotransposons^55^. Although infections have been linked to ibrutinib treatment, these are mostly bacterial and fungal infections^56,57^. However, there are reported cases showing the reactivation of hepatitis B virus (HBV) after ibrutinib treatment ^58,59^. Because APOBEC3G can also inhibit HBV, our finding may provide new insights to the understanding of HBV reactivations with ibrutinib treatment where imbalances or deficiencies of APOBEC3 family members contribute to a deficient host response to infections. In addition, if ibrutinib therapy is accompanied by other novel agents that modify the immune system or other mechanisms of host resistance to infections that could amplify this complication.

Recent work has found that APOBEC3 increases DNA replication stress and chromosome instability in early breast and lung cancer evolution^41^. Our data also implicated the role of APOBEC3 in the regulation of replication stress in malignant B cell lines. Although APOBEC3 have been implicated in the off-target mutations and evolution of CLL^60,61^, the functional importance of our finding in CLL is still limited. APOBEC3 are implicated in the generation of genomic mutations of various types of cancers^21^, thus we speculate that they can also drive gene mutations during the evolution of CLL. If true, our findings here would provide mechanistic insights that a signaling pathway regulated enhancer remodeling couples the extracellular environment to regulate the genetic evolution of leukemic cells. Indeed, our preliminary analysis shows that APOBEC3 are involved in the DNA replication stress in malignant B cells. We also found that enhancer regulated APOBEC3 expression is associated with DNA damage during S phase in the MEC1 cell line, which suggests that increased expression of APOBEC3 may induce transcription replication conflicts, a major driver of cancer evolution^62^. Future work will be focus role of the BCR regulated APOBEC3 expression in relation to alteration of immune resistance as it relates to infection propensity as well as their role in B cell genomic instability and CLL evolution.

## Acknowledgements

The authors thank all the members of the Mayo Stabile CLL Research group for helpful discussions of the work; and Dr. Rentian Wu for helpful insights and technique supports of the enhancer deletion assays; and Dr. Steven Offer for strategic discussions; and Qianqian Guo for help with the Edu/DAPI staining and flow cytometry analysis. We than Henry J. Predolin Foundation for the support for the Mayo Clinic CLL bank.

The work was supported by Hematology research merit grant, Mayo Clinic Cancer Center Hematological Malignancies Program, Hollis Brownstein Research Grants Program from Leukemia Research Foundation (Z.W.); R01 GM130588 from National Institutes of Health (C.Y.).

## Authorship

Contributions: Z.W. conceived and designed the project; Z.W., J.C.B., C.R.S., J.Z. performed the experiments; Z.W., H.Y., J.Z., and H.G. analyzed the data; E.B., C.Y., S.L.S., S.A.P., provides key resources and supervision; Z.W. and N.E.K supervised the project.

