## supplemental Files for "B Cell Receptor Signaling Drives APOBEC3 Expression Via Direct Enhancer Regulation in Chronic Lymphocytic Leukemia B Cells"

**Materials and methods**

| Antibodies | Vendor | Cat Num and Antibody ID |
|---|---|---|
| Mouse anti-NFATc1 (7A6) | Santa Cruz Biotechnology | Cat# sc-7294, RRID: AB_2152503 |
| Mouse anti-alpha-tubulin (IB) | Developmental Studies Hybridoma Bank | Cat#: 12G10, RRID: AB_1157911 |
| Rabbit polyclonal to Histone H3 | Abcam | Cat#: 12 ab1791, RRID: AB_302613 |
| Rabbit polyclonal to H3K4me1 | Abcam | Cat# ab8895, RRID: AB_306847 |
| Rabbit polyclonal to H3K4me3 | Abcam | Cat# ab8580, RRID: AB_306649 |
| Rabbit polyclonal to H3K27ac | Abcam | Cat#: ab4729, RRID: AB_2118291 |
| Mouse monoclonal ANTI-FLAG® M2 | Sigma-Aldrich | Cat#: F1804, RRID: AB_262044 |
| APOBEC3G (D9C6Z) Rabbit mAb | Cell Signaling Technology | Cat# 43584, RRID: AB_2799245 |
| Rabbit polyclonal to APOBEC3C | GeneTex | Cat# GTX102164, RRID: AB_2616091 |
| Rabbit polyclonal to 53BP1 | Novus | Cat# NB100-304, RRID: AB_10003037 |
| Anti-phospho-Histone H2A.X (Ser139) Antibody | Millipore | Cat# 05-636, RRID: AB_309864 |
| Phospho-Chk1 (Ser345) (133D3) Rabbit mAb antibody | Cell Signaling Technology | Cat# 2348, RRID: AB_331212 |
| Chk1 (2G1D5) Mouse mAb antibody | Cell Signaling Technology | Cat# 2360, RRID: AB_2080320 |
| Anti-RPA 32 kDa subunit Antibody (9H8) | Santa Cruz Biotechnology | Cat# sc-56770, RRID: AB_785534) |
| RDye® 800CW donkey anti-mouse IgG | LI-COR Biosciences | Cat#: 926-32212, RRID: AB_621847 |
| IRDye® 680RD donkey anti-rabbit IgG | LI-COR Biosciences | Cat#: 926-68073 RRID: AB_10954442 |
| Goat Anti-Mouse IgG (H+L) Antibody, Alexa Fluor 594 Conjugated | Thermo Fisher Scientific | Cat# A-11005, RRID: AB_141372 |
| Goat Anti-Rabbit IgG (H+L) Antibody, Alexa Fluor 488 Conjugated | Thermo Fisher Scientific | Cat# A-11008, RRID: AB_143165 |

**Cell culture**. All cell lines were maintained at 37°C in a humidified incubator with an atmosphere of 5% $CO_2$. HEK293T/c17 (HEK293T) cells were cultured in Dulbecco's Modified Eagle's Medium (DMEM; Corning, Corning, NY); MEC1 cells were cultured in RPMI-1640 medium (Corning). Media were supplemented with 10% fetal bovine serum (FBS; Invitrogen), 2 mM L-glutamate (Corning), and 1x penicillin/streptomycin solution (Invitrogen). MEC1 cells stably expressing Cas9 or dCas9-KRAB-MECP2 were cultured in MEC1 medium supplemented with 10µg/ml Blasticidin.

**CLL patients and the purification of their leukemic B-cells.** All patients studied for blood samples provided written informed consent according to the Declaration of Helsinki to the Mayo Clinic Institutional Review Board, which approved these studies. Informed consents were also obtained from healthy donors to obtain their bone biopsy samples. Primary CLL B cells were purified from blood samples of CLL patients using the RosetteSep B-cell enrichment kit (Stem Cell Technologies). The typical purification range of CD5+/CD19+ CLL B-cells was >95–99% as determined by flow cytometric analysis. CLL B-cells were cultured for optimum viability in serum-free AIM-V (Gibco) medium as previously described[1].

Quantitative RT-PCR (RT-qPCR). Total RNA was extracted using the Direct-zol RNA Kit (Zymo Research, Tustin, CA) following the manufacturer's recommendations. Reverse transcription into cDNA was performed using Superscript III and random hexamer primers (Life Technologies) according to the manufacturer's instructions. Quantitative PCR (qPCR) was carried out using iTaq Universal SYBR Green Supermix (Biorad) on a CFX96 Touch Deep Well Real-Time PCR System (Biorad) using primers listed below. Relative expression was calculated as 2^- (∆Ct (target)- ∆Ct (GAPDH)). Three independent experiments were performed; for each experiment, gene expression was assessed in triplicate.

**Western blotting**. Whole cell lysates were extracted in Laemmli buffer (60 mM Tris-Cl pH 6.8, 2% SDS, 10% glycerol, 5% β-mercaptoethanol, 0.01% bromophenol blue), separated by SDS-PAGE gel, and transferred to PDVF membrane (MilliporeSigma). Blots were blocked using Odyssey Blocking Buffer (LI-COR Biosciences, Lincoln, NE) prior to incubation with primary antibody at 4°C overnight. Secondary antibody incubations (anti-mouse, 1:5,000 dilution; anti-rabbit, 1:10,000 dilution; both LI-COR Biosciences) were performed for 1 hour at room temperature. Proteins of interest were visualized by Odyssey infrared imaging system (LI-COR Biosciences).

**Plasmids**. pLX-TRE-dCas9-KRAB-MeCP2-BSD was a gift from Andrea Califano (Addgene plasmid # 140690; http://n2t.net/addgene:140690; RRID: Addgene_140690). lentiGuide-Puro was a gift from Feng Zhang (Addgene plasmid # 52963; http://n2t.net/addgene:52963; RRID: Addgene_52963). lentiCas9-Blast was a gift from Feng Zhang (Addgene plasmid # 52962; http://n2t.net/addgene:52962; RRID:Addgene_52962).

**Lentivirus production**. Lentiviral plasmids were co-transfected into HEK293T cells with packaging vectors psPAX2 (Addgene plasmid #12260; a gift from Didier Trono) and pMD2.G (Addgene plasmid #12259; a gift from Didier Trono) using PEI 4000 (Polysciences). Virus-containing medium was collected 48 hours after transfection and cleared of potential cells using 0.45-μm Steriflip filter units (MilliporeSigma).

**CRISPR-cas9 knockout**. Single guide RNAs (sgRNAs) targeting NFATc1 were designed using the Broad Institute Genetic Perturbation Platform (http://portals.broadinstitute.org/gpp/public/analysis-tools/sgrna-design) and cloned to lentiCRISPR V2. Lentivirus was packaged as described above, mixed with polybrene (final concentration 8 μg/mL; MilliporeSigma), and transduced to MEC1 cells. The APOBEC3 enhancer deletions were generated using CRISPR-Cas9 methods. Cas9 expressing MEC1 cells were generated with the infection of the MEC1 cells with lentiCas9-Blast and the cells were cultured with 10 μg/ml Blasticidin. Single guide RNAs (sgRNAs) were designed using the Broad Institute Genetic Perturbation Platform (http://portals.broadinstitute.org/gpp/public/analysis-tools/sgrna-design). Oligonucleotides were synthesized by Integrated DNA Technologies (IDT, Coralville, IA) and cloned into the lentiGuide-Puro plasmid. Lentivirus was packaged as described above, mixed with polybrene (final concentration 8 μg/mL; MilliporeSigma), and used to transduce MEC1 Cas9 cells. Twenty-four hours after infection, stable transduced cells were selected for 7 days in 0.5 μg/mL puromycin-containing medium. The deletion of the enhancers was determined by PCR and changes in APOBEC3 transcript levels were monitored using RT-qPCR.

**CRISPR-dCas9-Krab-MECP2 inactivation.** gRNAs targeting AEs were cloned to lentiGuide-Puro. MEC1 cells were infected with pLX-TRE-dCas9-KRAB-MeCP2-BSD and selected with 10 μg/ml Blasticidin, then the pool populations were infected with two gRNAs targeting enhancer regions and selected with Puromycin. Two days after selection and the cells were incubated with doxycycline (0.5 μg/ml) for 5 days then harvested for analysis.

**CUT&Tag.** CUT&Tag was performed as described (https://www.protocols.io/view/bench-top-cut-amp-tag-bcuhiwt6/abstract). CUT&Tag libraries were sequenced to 50 base pairs on an Illumina HiSeq 4000 using pair-end mode at the Mayo Clinic Gene Analysis Shared Resources.

**CUT&Tag data analysis**. CUT&Tag data were analyzed following CUT&Tag Data Processing and Analysis Tutorial (https://www.protocols.io/view/cut-amp-tag-data-processing-and-analysis-tutorial-bjk2kkye). Reads were aligned to the human genome (hg38) or E. coli genome using Bowtie2 [2]. The fragments mapped to E. coli were used as the Spike-in calibration. Uniquely mapped reads were used for further analysis. Peaks were identified using MACS2 [3] using 0.01 as the cutoff FDR value. Mapped reads were transformed to BigWig files by deepTools [4] for visualization. Genome-wide coverage and read density scan were calculated by ngs.plot [5] using Reads Per Kilobase per Million mapped reads (RPKM) for normalization. Gene ontology analysis was performed by GREAT v3.0.0 [6].

**RNA-seq.** Total RNA was extract using the Direct-zol RNA Kit (Zymo Research). Library preparation and sequencing were performed using the NovaSeq 6000 platform, paired-end 150 bp by Novogene (Sacramento, CA).

**RNA-seq data analysis**. Reads were aligned using Kallisto[7]. Transcript abundance files were then used in the DESeq2 R package, which was used for all downstream differential expression analysis and generation of volcano plots. Differentially expressed genes between samples were compared with a cutoff of log2 Fold Change > 0.5, $p<0.05$.

**ATAC-seq**. ATAC-seq library construction was performed as previously described [8]. Fifty thousand cells were lysed in cold ATAC-Resuspension Buffer (RSB) containing 0.1% NP40, 0.1% Tween 20, and 0.01% digitonin on ice. Lysis buffer was washed out with cold ATAC-RSB containing 0.1% Tween 20 followed by centrifugation at 4°C. Pellets (nuclei) were resuspended in transposition mix containing Tagment DNA buffer, Tn5 Transposase, and 0.05% Tween 20. Reactions were incubated for 30 min at 37°C with constant agitation. Transposed DNA was purified using QIAgen MinElute columns. DNA was amplified using Nextera sequencing primers (Illumina) and NEB High Fidelity 2X PCR Master Mix (New England Biolabs) for 3–5 cycles. PCR-amplified DNA was purified using QIAgen MinElute columns and sequenced using an Illumina HiSeq 4000 with single-end reads of 50 bases by the Mayo Clinic Gene Analysis Shared Resources.

**ATAC-seq data analysis.** ATAC-seq data analysis was performed as previously reported [8,9]. The following tools were used: Samtools, Picard, Bowtie2, macs2, and bedtools v2.23.0.

Nextera adaptor sequences were first trimmed from the reads by Trim Galore!. These reads were aligned to a reference genome using bowtie2 with standard parameters and a maximum fragment length of 2,000. Picard was used to remove duplicate reads. De-duplicated reads were filtered for high quality (MAPQ ≥ 30), nonmitochondrial, non-Y chromosome, and properly paired (Samtools flag 0 × 2) reads.

**Statistical analysis**. Significance was defined as $p \leq 0.05$ unless otherwise noted in the text. Paired or un-paired two-tailed Student's t test and two-way ANOVA statistics were calculated using GraphPad Prism 9. Two-tailed Student's t tests were used for pairwise comparisons as noted in the text.

**Supplementary Fig.1**

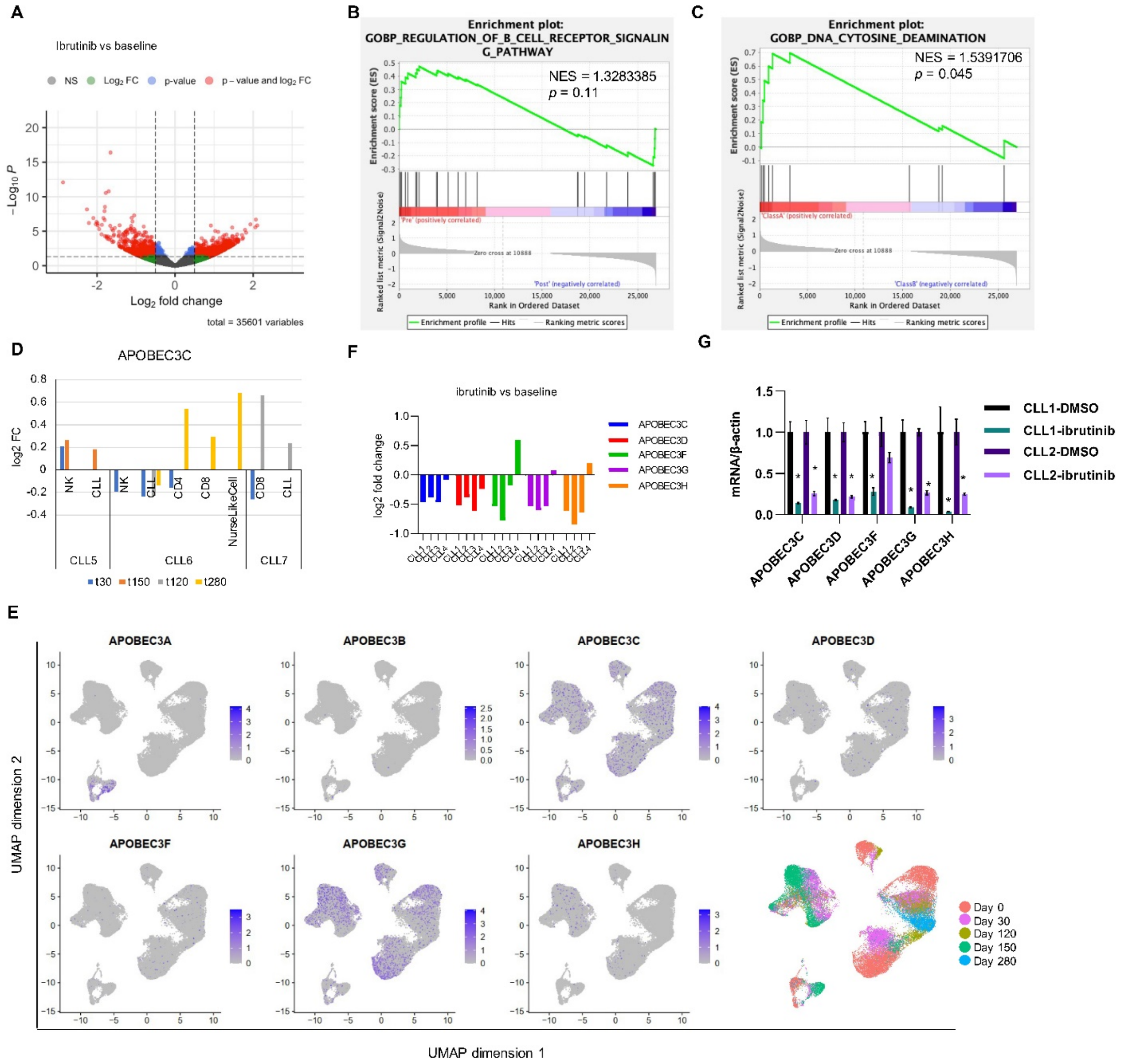


### Supplementary Fig. 1 Ibrutinib treatment induces genes expression changes in CLL B cells.

A) Differentially expressed genes in CLL B cells from one-year of continues ibrutinib treated patients compared to that from pretreated patients.

B) Gene set enrichment analysis showing enrichment of BCR signaling pathway signatures in ibrutinib pretreated CLL cells compared to CLL cells from patients with one-year of continuous ibrutinib treatment.

C) Gene set enrichment analysis showing enrichment of DNA cytosine deamination gene signatures in CLL B cells from patients before ibrutinib treatment compared to that from patients after one-year of continuous ibrutinib treatment.

D) APOBEC3C expression in indicated cells from ibrutinib treated patients. t30 means 30 days after initiation of ibrutinib treatment.

E) Feature plots showing the expression of APOBEC3 genes.

F) RT-qPCR analysis of APOBEC3 expression of CLL B cells from patients pre and after one-year ibrutinib treatment. baseline = pre ibrutinib treatment; ibrutinib = one-year of continuous ibrutinib treatment.

G) RT-qPCR analysis of APOBEC3 in CLL B cells treated with 2.5μM ibrutinib for 3 days. n = 3 independent experiments for the CLL B cells from each patient.

**Supplementary Fig. 2**

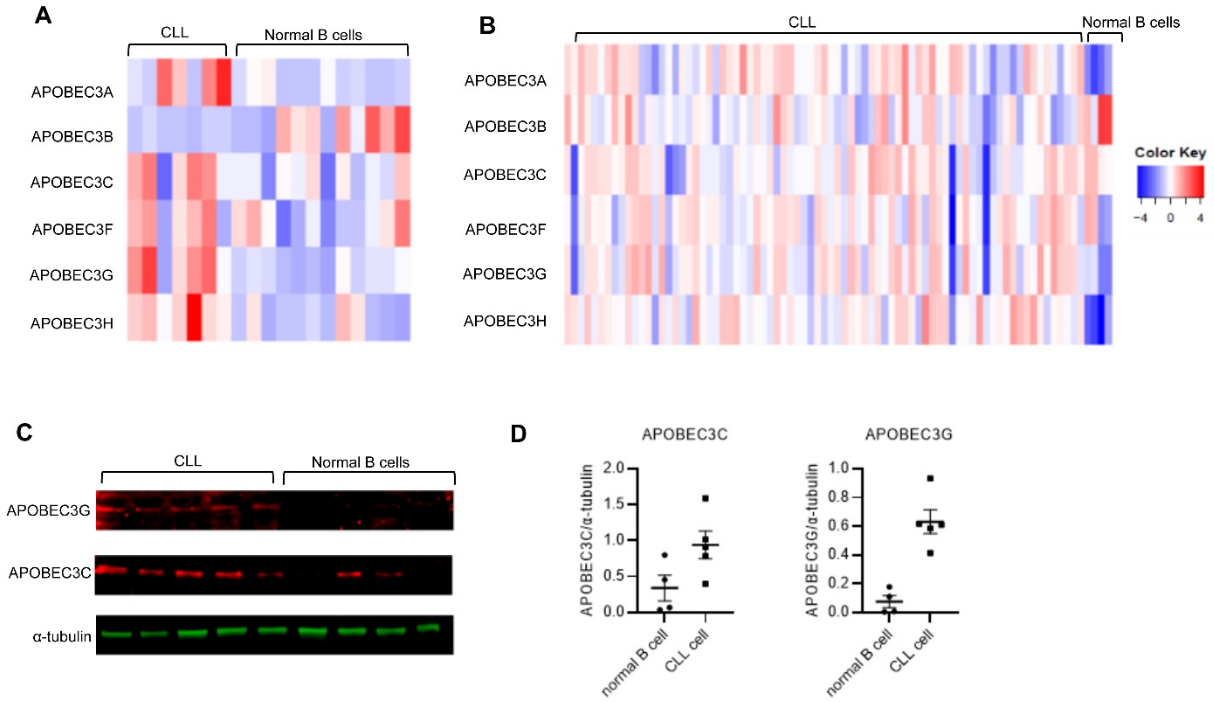


**Supplementary Fig. 2 APOBEC3s expression in normal and CLL B cells.**

A) and B), Heatmaps showing the relative expression of APOBEC3s in normal and CLL B cells. RNA-seq datasets were obtained from EGAD00001004046 (A) and GSE119103 (B).

C) Western blot showing APOBEC3C and APOBEC3G level in CLL B cells and normal B cells.

Supplementary Fig. 3

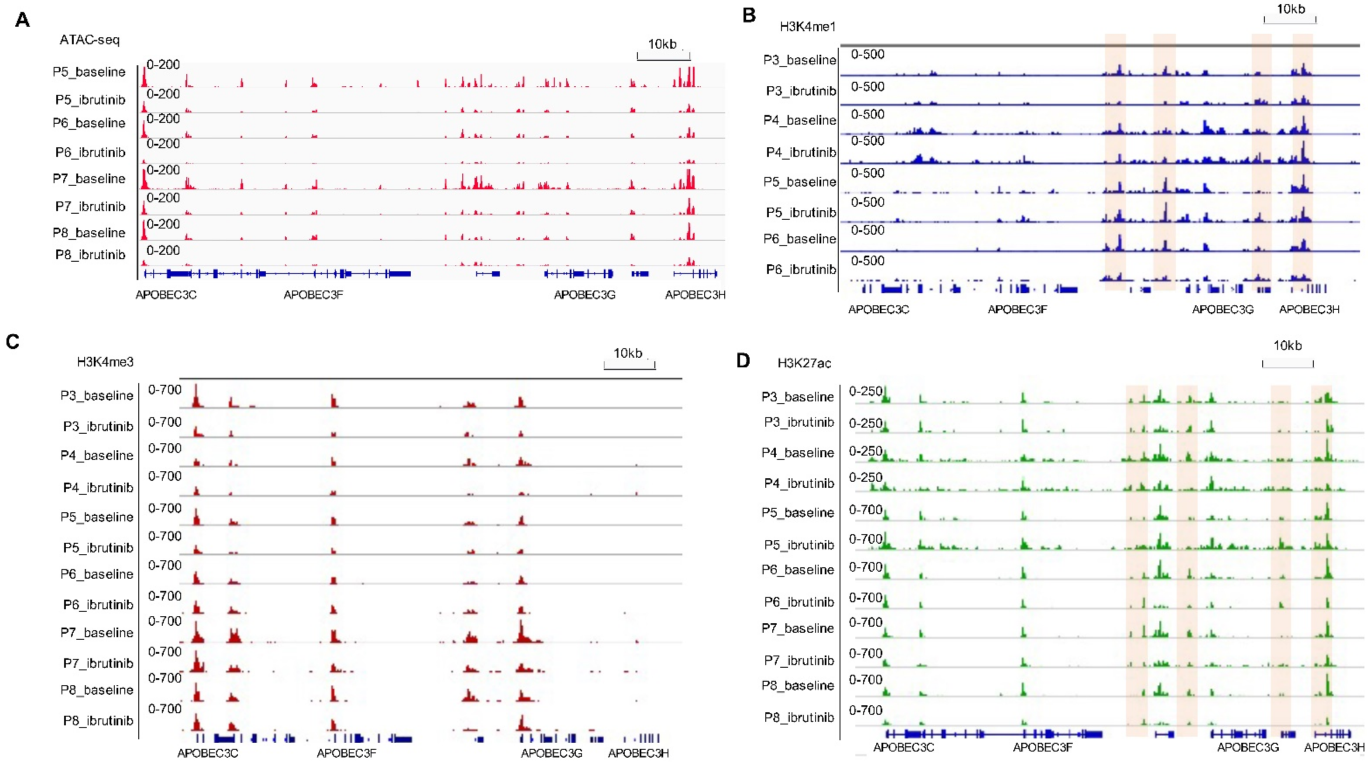


**Supplementary Fig. 3 Ibrutinib treatment leads to decreased Chromatin accessibility, H3K4me1, and H3K27ac at APOBEC3 enhancers in CLL cells.**

A-D) Genome tracks showing ATAC-seq, CUT&Tag of H3K4me1, H3K4me3, and H3K27ac profiles of putative APOBEC3 enhancers. Light brown shadows show enhancers. P, patient; baseline = pre ibrutinib treatment; ibrutinib = one-year of continuous ibrutinib treatment.

### Supplementary Fig. 4

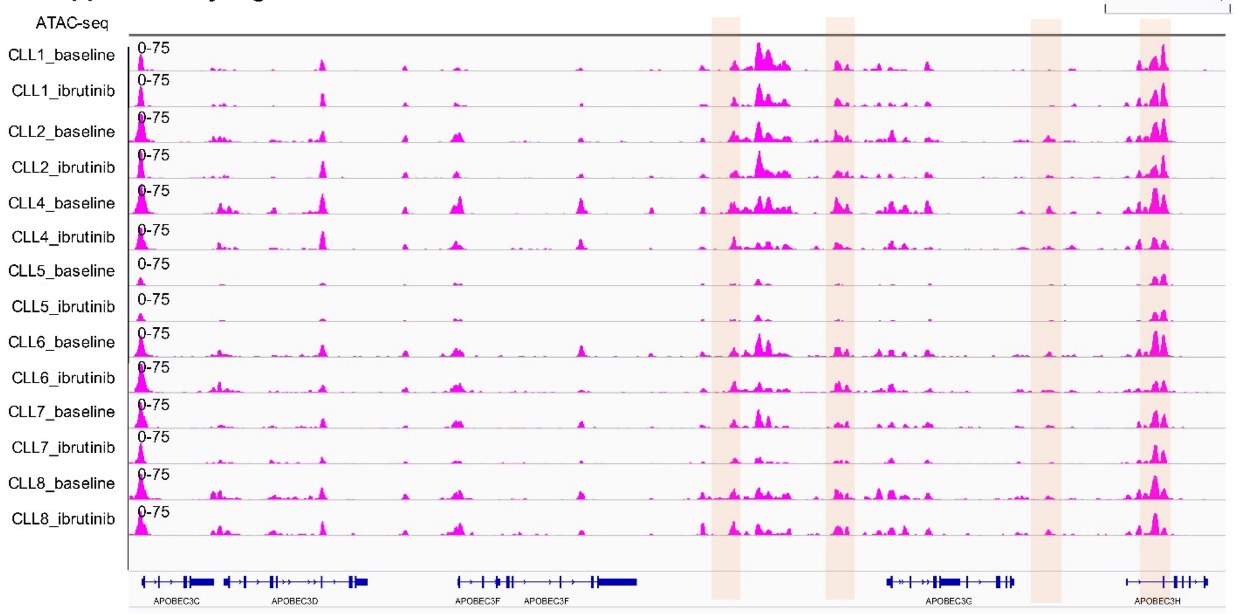


**Supplementary Fig. 4 Ibrutinib treatment leads to decreased chromatin accessibility at APOBEC3 enhancers.**

Genome tracks showing ATAC-seq profiles of putative APOBEC3 enhancers. P, patient; baseline = pre ibrutinib treatment; ibrutinib = 120 days post initiation of ibrutinib treatment. Data are downloaded from GSE111015.

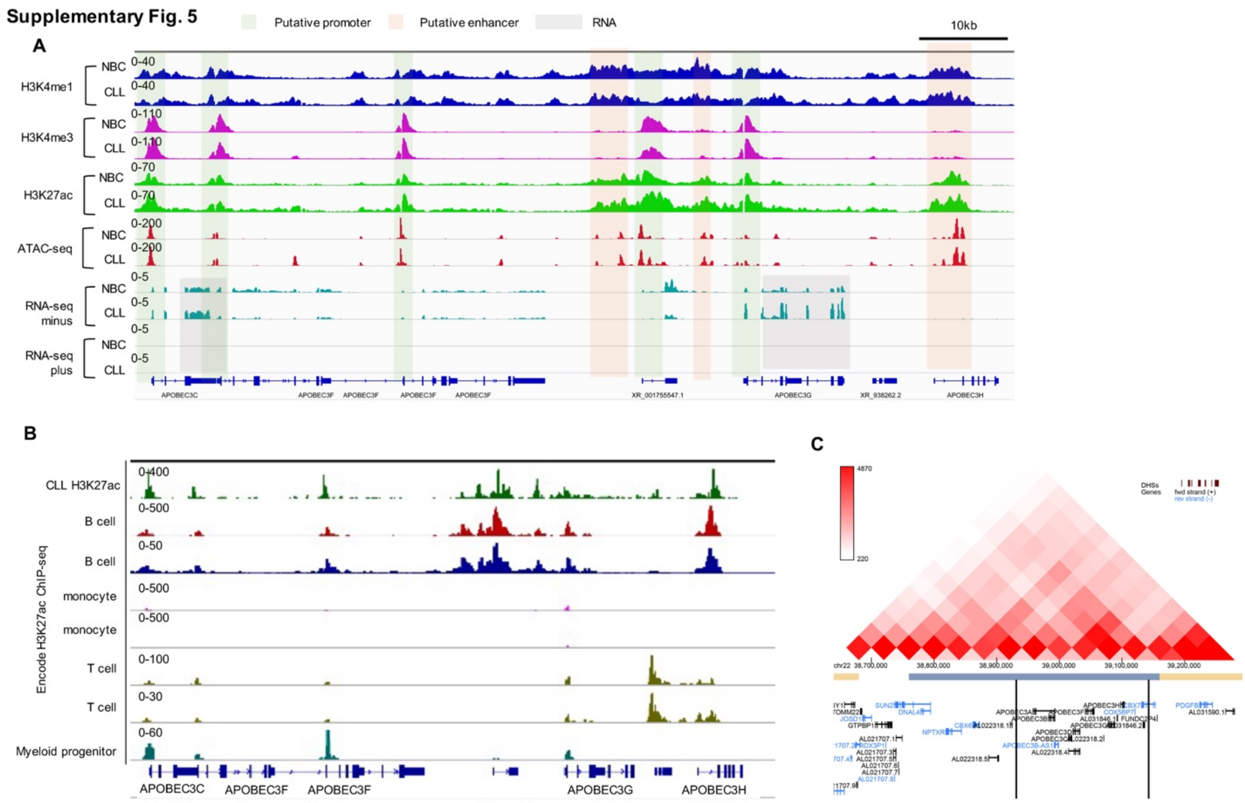


**Supplementary Fig. 5 The putative APOBEC3s enhancer signatures are B cell specific.**

A) Genome tracks showing H3K4me1, H3K4me3, H3K27ac profiles and ATAC-seq and RNA-seq of putative APOBEC3 enhancers in normal and CLL B cells. Data were from downloaded from published dataset EGAD00001004046[10].

B) Genome tracks showing H3K27ac in indicated cells. ChIP-seq data were downloaded from ENCODE project[11].

C) The Hi-C analysis from GM12878 cells is analyzed and taken from the 3D genome browser site[12]. The boundaries of the topologically associated domains (TADs) and the location of APOBEC3s is indicated below.

**Supplementary Fig. 6**

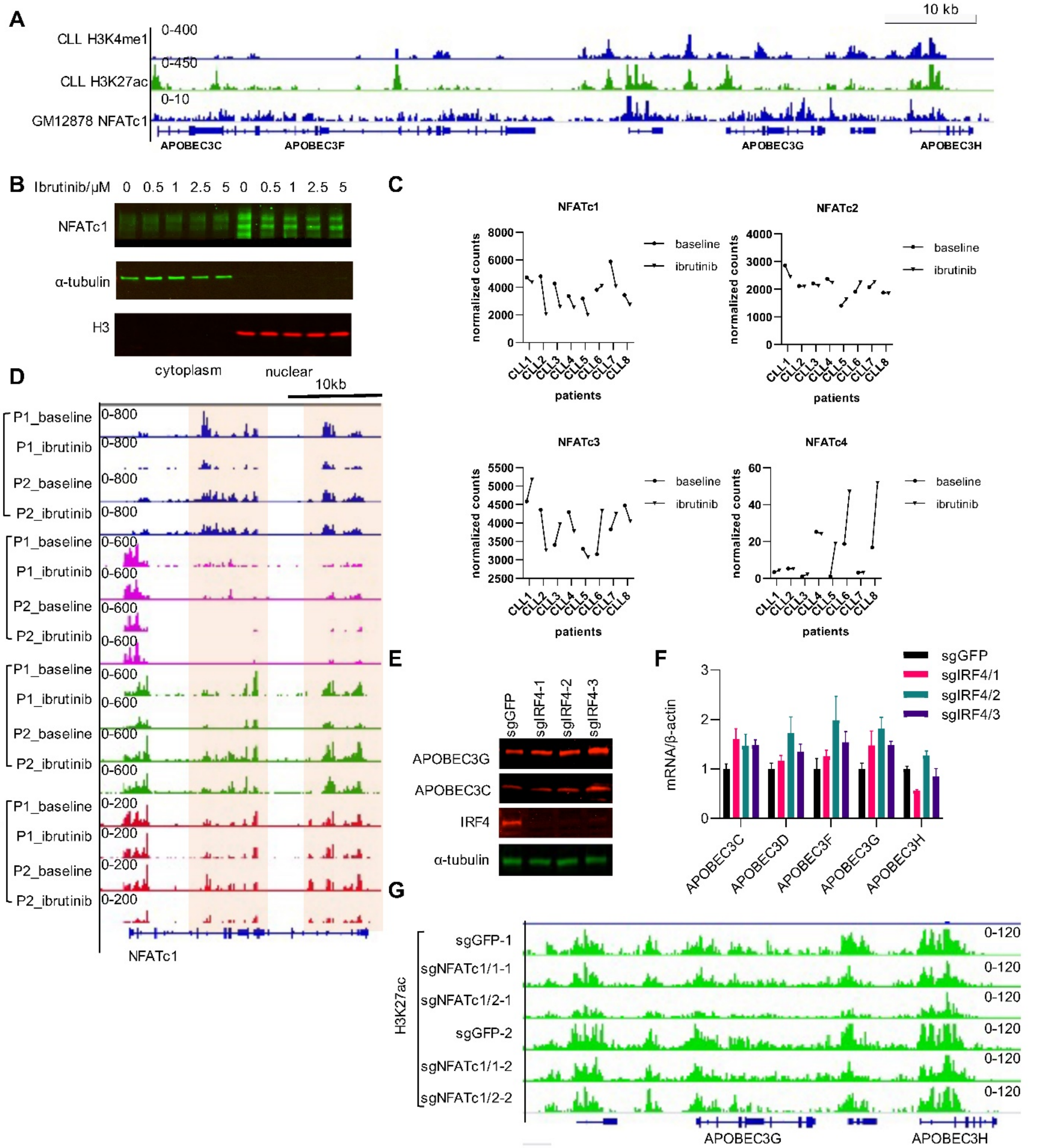


**Supplementary Fig. 6 NFATc1 controls the APOBEC3 enhancers activity.**

A) Genome tracks showing H3K4me1, H3K27ac profiles in CLL B cells and NFATc1 profile in GM12878 cells. NFATc1 ChIP-seq data were downloaded from ENCODE project[11].

B) Western blot showing the NFATc1 level in nuclear and cytoplasm of CLL B cells treated with ibrutinib as indicated for 24 hours.

C) NFATc1-4 expression defined by RNA-seq in CLL B cells from patients before and after ibrutinib treatment.

D) Genome tracks showing CUT&Tag of H3K4me1, H3K4me3, H3K27ac and ATAC-seq profile around NFATc1.

E) Western blot analysis of APOBEC3 expression in IRF4 depleted MEC1 cells.

F) RT-PCR analysis of APOBEC3 expression in IRF4 depleted MEC1 cells.

G) Genome tracks showing H3K27ac profiles at APOBEC3 enhancers from CUT&Tag. Note: This snapshot is with a different scale from the Fig. 4G.
